# A rapid, cost-effective, colorimetric LAMP assay (CLASS) for detecting invasive malaria vector, *Anopheles stephensi*

**DOI:** 10.1101/2024.02.06.579110

**Authors:** Cristina Rafferty, Gloria Raise, Jenyiah Scaife, Bernard Abongo, Seline Omondi, Sylvia Milanoi, Margaret Muchoki, Brenda Onyango, Eric Ochomo, Sarah Zohdy

## Abstract

*Anopheles stephensi*, an invasive malaria vector in Africa, has the potential to impact the landscape of malaria on the continent, threatening to put an additional 126 million people per year at risk of malaria, largely in peri-urban/urban areas. To accelerate the early detection and rapid response to *An. stephensi* and ensure no gains made in malaria control and elimination are lost, it is critical to confirm the presence of the species and the geographic extent of its spread to inform control. However, morphological identification may be misinterpreted if specimens are damaged and existing molecular species confirmation assays require specialized laboratory equipment and training and may be challenging to interpret, requiring additional sequencing confirmation. A colorimetric rapid loop-mediated isothermal amplification (LAMP) assay for molecular *An. stephensi* species identification was developed and optimized. The colorimetric assay requires only a heat source and reagents and can be used with or without DNA extraction resulting in positive color change in 30-35 minutes. To determine analytical sensitivity, a 1:10 dilution series of the DNA extract was conducted showing 100% assay sensitivity down to 0.003 nanograms. To determine specificity, three different *An. stephensi* laboratory strains (STE2, SDA 500, UCI), 8 other Anopheles mosquito species, and *Aedes aegypti* were compared, and the results indicated 100% specificity across these species. To determine use without the need for DNA extraction, samples evaluated included a single mosquito leg, whole adult or larval mosquitoes, and pooled DNA extract from several mosquito species. A total of 1687 individual reactions were tested during optimization and all LAMP assay results were compared against the conventional PCR assay and confirmed through Sanger sequencing. To validate the optimized assay on wild caught specimens, DNA extracted from 12 wild caught, sequence-confirmed *An. stephensi* from Marsabit, Kenya, were tested and the colorimetric assay was accurate in identifying all of the specimens as *An. stephensi*. The assay described presents an opportunity to accelerate *An. stephensi* molecular identification in new and existing locations in Africa, within its endemic range, and globally. These findings present a simple, rapid, unique alternative to existing PCR and sequencing-based *An. stephensi* species identification and confirmation strategies. With additional field validation studies, molecular screening tools like the colorimetric LAMP-based An. *stephensi* species identification (CLASS) assay fill an important gap of rapid confirmation of this invasive vector and presents an ideal opportunity to better understand the spread of the species in Africa and other recently invaded areas, thus accelerating a response to mitigate its long-term impacts on malaria on the continent.

## Introduction

In 2012, *Anopheles stephensi*, a primary malaria vector in south Asia and the Arabian Peninsula, was detected in Djibouti, a country that was approaching malaria pre-elimination status in 2011. Unlike most typical African malaria vectors, *An. stephensi* has an ability to thrive in both urban and rural environments. Following the detection in Djibouti, *An. stephensi* was reported in Ethiopia and Sudan in 2016, Somalia in 2019, Nigeria in 2020, Kenya in 2022, and Ghana and Eritrea in 2023 (WHO, 2023). Following the initial detection in Djibouti, which was found in association with a malaria outbreak (Faulde et al., 2014), a 36-fold increase in malaria was reported in the country from <2,000 cases per year in 2012 to >73,000 cases in 2020 (WHO, 2021). Similarly, in 2022 in Dire Dawa, the second largest city in Ethiopia, an unusual dry season outbreak of malaria was reported and follow on epidemiological and entomological investigations incriminated the role of *An. stephensi* in driving this urban malaria outbreak (Emiru et al. 2023). Furthermore, its insecticide resistance and biting and resting trends present a challenge to current proven malaria vector control tools, such as insecticide treated bed nets (ITNs) and indoor residual spraying (IRS) (Faulde et al., 2016; Balkew et al. 2021). Modeling studies have predicted that if *An. stephensi* continues to spread throughout Africa, an additional 126 million people, predominantly in urban areas, will be at risk of malaria each year (Sinka et al, 2020; Hamlet et al, 2022). To respond to this threat, WHO has launched an initiative to halt the spread of *An. stephensi* (WHO, 2022), and the U.S. President’s Malaria Initiative released an action plan (PMI, 2021 and 2023) to encourage enhanced surveillance for the species to allow for early detection in new locations and rapid response to halt spread and mitigate impacts. Early detection of the species through enhanced surveillance is essential to determine the current extent of the species on the continent to establish mitigation, containment, and even elimination strategies.

Despite efforts to enhance surveillance for *An. stephensi* in Africa, including a revised WHO vector alert and guidance (WHO, 2023), being an invasive species not historically included in morphological keys until 2020 (Coetzee, 2020), the vector may be missed in routine malaria surveillance activities and follow up molecular identification without specific optimized tools. For example, *An. stephensi* may be misidentified as the common malaria vector *An. gambiae* s.l. if training on morphological identification is inadequate (WHO, 2019). Additionally, reporting a detection of *An. stephensi* in a new country to WHO requires molecular confirmation, which may be challenging in resource limited settings. *Anopheles stephensi* surveillance often requires larval surveys (Ahmed et al. 2021) because adult collections routinely used in malaria vector surveillance are not optimal for the species (Balkew et al. 2021). Larval samples must be reared to adults for morphological identification, as there is not currently a validated morphological key to identify *An. stephensi* larvae to the species level, so larval samples that do not emerge to adults may also require molecular confirmation.

In 2023, a new PCR protocol for *An. stephensi* species identification was released (Singh et al., 2023) which was shown to detect *An. stephensi* even among pooled samples and provides a promising avenue for *An. stephensi* detection close to the site of collection. However, PCR processing of suspect specimens may be time consuming and limited by molecular laboratory capacity, access to reagents, trained personnel, and assay specificity and interpretation.

Loop-mediated isothermal amplification (LAMP) assays have been used since the 1990s for rapid amplification of gene targets (Garg et al., 2022). The LAMP assays function like PCR amplifying targeted genes resulting in a visual change through fluorescence, turbidity, or color of amplified products, providing a qualitative indicator of positivity. In this way, LAMP assays function like conventional PCR assays which yield a band (positive) or no band (negative). However, LAMP assays do not require temperature cycling for denaturing and annealing like conventional PCR, and instead produce copies through multiple looped primer sets at one consistent heating temperature, removing the need for a thermal cycler and instead requiring only a heat block, water bath, or any other tool or device that keeps temperature consistent for a period of time. In the literature, one study even used hand-warmers and a Styrofoam cup to conduct a LAMP assay (Hatano et al. 2010). LAMP technology has evolved over the years and with the COVID-19 pandemic’s need for rapid diagnostics this technology expanded to include colorimetric and dipstick assays.

To address the challenges that invasive *An. stephensi* surveillance and accompanying molecular confirmation presents, the aim of the study was to develop an easy-to-interpret, rapid colorimetric LAMP-based *Anopheles stephensi* species identification assay, from here on referred to as CLASS assay, specifically designed and optimized for molecular species identification of *An. stephensi*, for use in resource limited settings or for rapid high-throughput screening. To ensure accuracy and feasibility for deployment of the developed assay, the following objectives were included: 1) design of optimal primers and assay conditions, 2) determination of assay sensitivity and pooling strategies, 3) determination of assay specificity when compared to congeners or conspecifics, 4) development of direct sample amplification approaches without the need for DNA extraction, 5) comparison of results between the published PCR protocol and CLASS assay, and 6) evaluation of CLASS on wild caught, sequence confirmed, invasive *An. stephensi* collected from Kenya.

## Methods

### LAMP primer design and optimization

LAMP primers for this study were designed using the New England Biolabs, Inc. NEB®LAMP Primer Design Tool version 1.4.1 (https://lamp.neb.com) based on the internal transcribed spacer 2 (ITS2) rDNA region unique to *An. stephensi* using the sequence from GenBank accession number MW732931.1. Attempts to set fixed primers resulted in no possible loop primer combinations by the program, therefore default parameters for the tool were used and the program was allowed to choose primers on its own. Of four possible primer sets tested, two contained primers in the species-specific region and were tested across a temperature gradient, with two sets of differing concentrations. Initial test concentrations were adapted from NEB kit manufacturer recommendations, and primer concentrations used in the published *An. gambiae* species ID LAMP assay publication (Bonizzoni et al, 2009). One primer set showed positive, consistent results and minimum cross-reactivity to other species. This primer set was then optimized further for maximum specificity (Figure 1).

**Figure 1.**
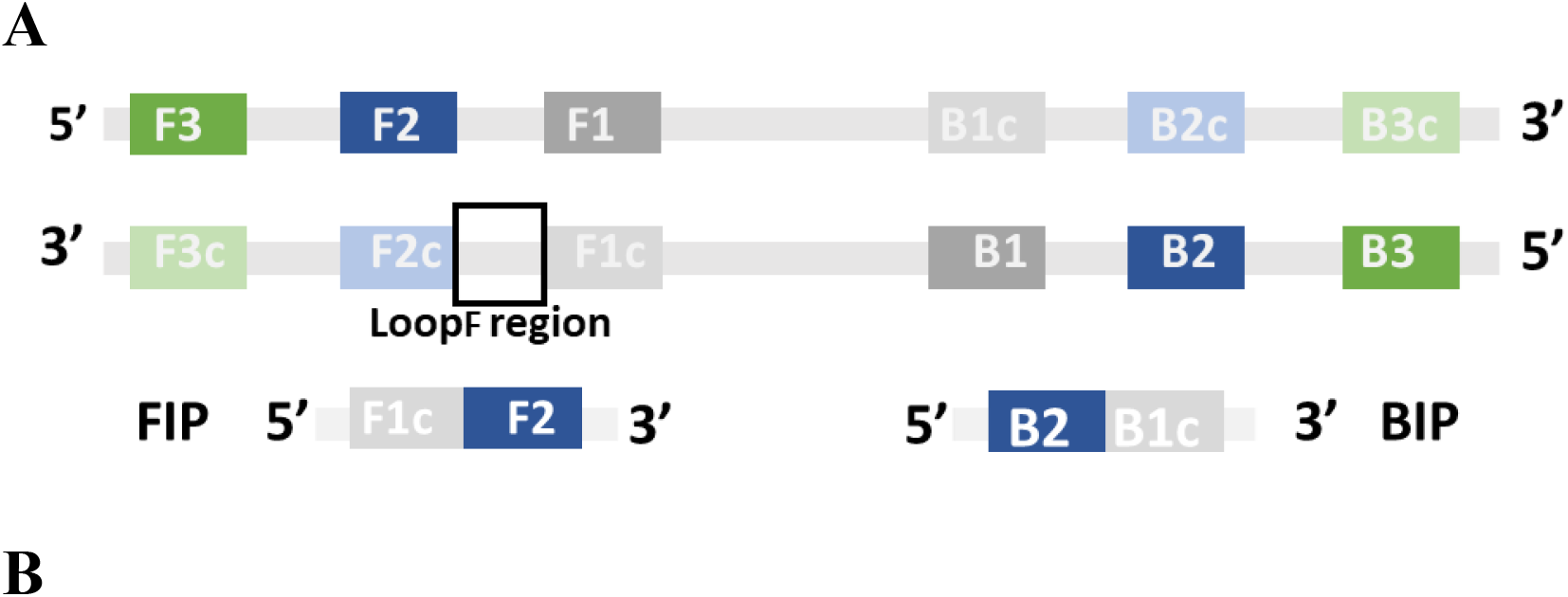
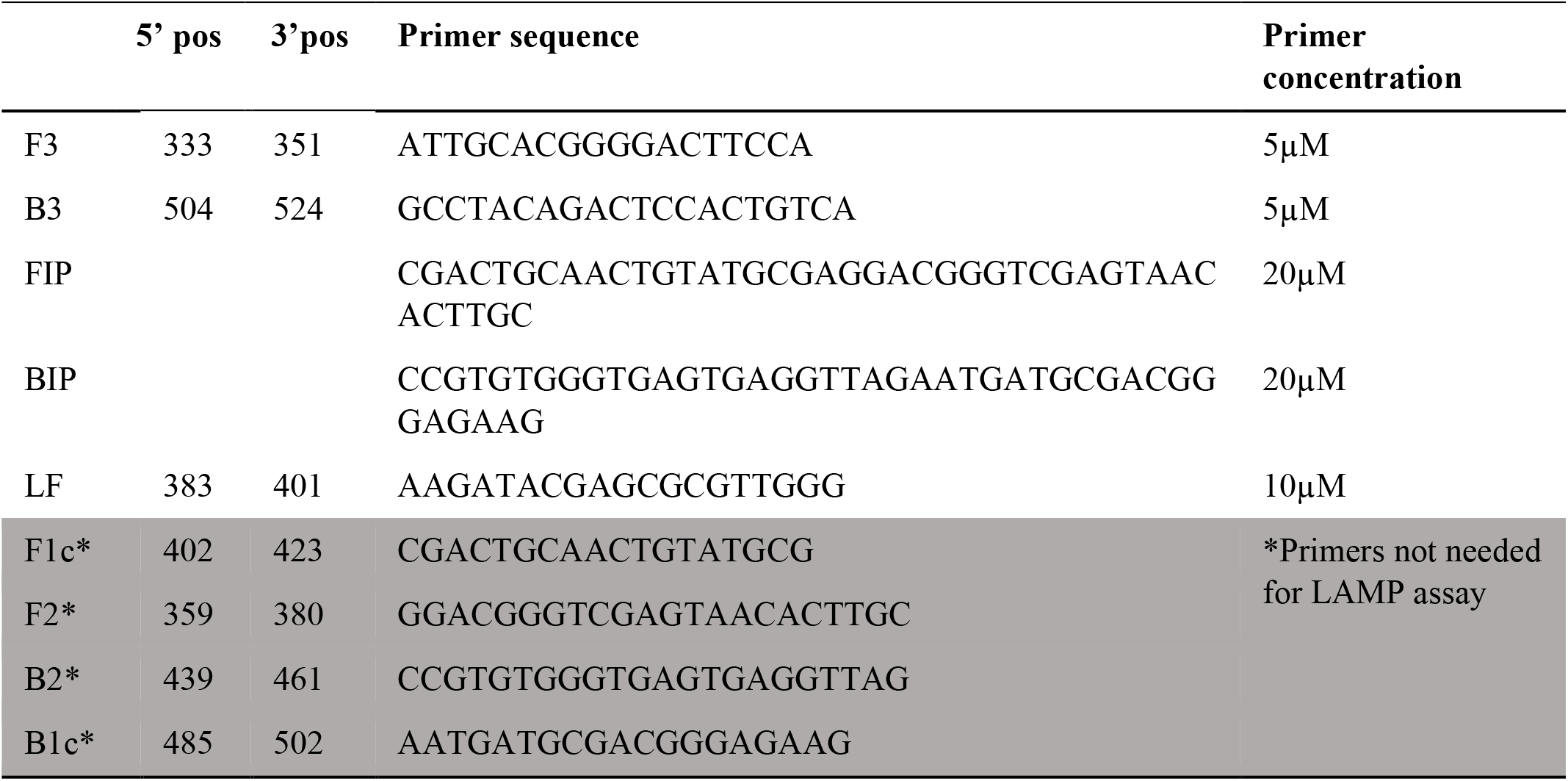
Looped primers were designed using the NEB LAMP Primer Design Tool targeting *An. stephensi* ITS2 sequence regions. **A**. A schematic of the primer design process ensuring looping to amplify at an isothermal temperature was generated and **B**. The five primers F3, B3, FIP, BIP, and LF selected for the colorimetric LAMP-based *Anopheles stephensi* species identification (CLASS) assay and identified concentrations for optimized specificity and sensitivity.

### Insectary reared An. stephensi and other mosquito species

Adult insectary-reared and maintained colony mosquitoes from eight distinct non-*An. stephensi Anopheles* species, three strains of *An. stephensi* of different origins (STE2, SDA 500, UCI), and one *Aedes* (*Aedes aegypti*), were obtained from the Malaria Research and Reference Reagent Resource Center (MR4) through BEI resources (Table 1, Acknowledgements).

**Table 1.** Mosquito species used in this study and their corresponding strains and catalog numbers from BEI resources.

| Species | MRA (BEI reference) | Strain name |
| --- | --- | --- |
| *An. gambiae* s.s. | 112 | G3 |
| *An. gambiae* s.s. / *An. coluzzii* hybrid | 334 | RSP |
| *An. coluzzii* | 1280 | AKDR |
| *An. arabiensis* | 856 | DONGOLA |
| *An. stephensi* | 1323 | UCI |
| *An. stephensi* | N/A | SDA500 |
| *An. funestus* s.s | N/A | FANG |
| *An. quadriannulatus* | 1155 | SANGWE |
| *An. dirus* | 700 | WRAIR |
| *An. merus* | 1156 | MAF |
| *An. stephensi* | 128 | STE2 |
| *Aedes aegypti* | 734 | ROCK |

### DNA extraction

DNA from either whole adult, a single mosquito leg, or a whole 3^rd^ instar larva was extracted using the Extracta DNA Prep for PCR ®kit (Quantabio Beverly, MA) adapted as follows: mosquito material was added to a PCR tube containing either 25µl (for a single leg) or 50µl (for a whole adult or larva) of Quantabio Extraction Reagent® and incubated in a thermal cycler at 95°C for 30 minutes. An equal volume of Quantabio Stabilization Buffer® was added, and DNA was stored at -20°C until further analysis. For the pooled species sample, 1µl of DNA from 9 DNA extractions of non-*An. stephensi* mosquito species, and 1µl of DNA extracted from *An. stephensi* (STE2) were combined in a microfuge tube and contents mixed.

### *Colorimetric LAMP-based* An. stephensi *identification (CLASS) assay*

CLASS reactions were carried out using the NEB WarmStart^®^Colorimetric LAMP 2X Master Mix, following manufacturer recommendations but optimized as follows: 1µl of genomic DNA was added to a tube containing 12.5µl of WarmStart^®^Colorimetric LAMP 2X Master Mix, 10X primers at the following concentrations: 5µM of B3 and F3 primers, 20µM of BIP and FIP primers, 10µM of LF primer. Molecular grade water was added to reach a final volume of 25µl. Reaction tubes were placed in a thermal cycler at 65°C for 30 minutes and inspected visually for color change. A positive amplification appears yellow, whereas a negative result remains pink. Primers were tested on extracted DNA from 12 assorted insectary-reared adult and larvae, including three *An. stephensi* strains (STE2, SDA500, and UCI), 14 field-collected specimens, including three sequence-confirmed *An. stephensi*, and DNA from pooled species. A no DNA control was included in each run of the assay.

### Analytical sensitivity determination

To test the sensitivity of the assay, a serial dilution (1:10) of DNA extract from a whole mosquito from the UCI *An. stephensi* strain was made. Starting DNA concentration was determined by using a Thermo Scientific NanoDrop 2000c spectrophotometer on 1uL of DNA extract. The extract was then used to create a 1:10 dilution series, testing each concentration on the CLASS assay to determine whether a positive color change would result. A negative result was determined with no color change (remained pink), whereas a positive result was determined with a full color change to yellow. Samples from the dilution series were run in triplicate until full color change was no longer detected; two additional dilutions were run to determine the sensitivity cut off.

### Specificity determination

Optimized primers were tested against 12 laboratory anopheline strains which included three strains of *An. stephensi* (SDA500, STE2, UCI) and one strain of *Aedes aegypti* (Table 1). Additionally, the assay was subsequently tested against 96 individual mosquitoes from each *An. stephensi* laboratory strain and 48 *An. gambiae, An. coluzzii, An. arabiensis, An. funestus*, and *Aedes aegypti* laboratory-reared samples for specificity and cross reactivity. All reactions were run in triplicate to generate data on cross-reactivity of the assay with other species and the specificity of the assay to *An. stephensi*. Three strains of *An. stephensi* were included to determine if there is any variation in target specificity across *An. stephensi* of different origins. As a qualitative colorimetric assay, a full color change from pink to yellow indicated a positive test result and samples which remained bright pink or lighter pink were designated negative.

### Sample use for LAMP amplification

To determine whether DNA extract is needed to run the CLASS assay or if tissue (mosquito leg, full larval, full adult mosquito) or pooled DNA amplified successfully, single insectary-reared mosquito legs were inserted directly into the master mix and compared to DNA extracted from a single leg. Because *An. stephensi* are often collected as larvae, the assay was also tested by immersing a whole larva into the master mix and using extracted DNA from a single larva. Partial testing of eDNA using larval pan water in lieu of extracted DNA was also carried out using 1 µl of larval pan water in lieu of DNA. The assay was also tested against whole adult mosquitoes and compared to whole adult mosquito DNA. Additionally, pooled DNA extract from whole adult mosquitoes and from individual legs from nine mosquito strains and one *An. stephensi* strain was also tested.

### Conventional PCR comparison

To determine ease of interpretation of the CLASS assay, LAMP results were compared to results from the conventional *An. stephensi* species identification PCR assay (Singh et al., 2023) following the publication conditions and adapted as follows: 2X Quantabio Accustart® PCR mix, 10µM of each primer, molecular water to reach a final volume of 20µl, and using 1µl of the extracted DNA from same species used in the CLASS assay. A successful PCR reaction yields a control band of varying sizes for all mosquito species and an additional *An. stephensi* specific band of 438bp for a positive *An. stephensi* sample.

### CLASS assay validation on wild-caught invasive An. stephensi *from Kenya*

Sequence confirmed DNA extract from wild-caught invasive *An. stephensi* from Kenya (GenBank Accession numbers OQ275144, OQ275145, OQ275146, OQ878216, OQ878217, and OQ878218) were also run using the CLASS assay to determine whether a positive color change would result from wild caught mosquito DNA extract that was sequence confirmed previously (Ochomo et al. 2023).

Additionally, a set of 55 wild-caught samples from Marsabit, Kenya which previously failed to amplify via conventional PCR assay, and had been previously sequenced with COI barcoding primers for species identification (Folmer et al, 1994) were tested using the CLASS assay. DNA extracted at the KEMRI lab was dried and shipped to CDC. Samples were resuspended in 25µL of PCR-grade water and stored at - 20°C until processed.

## Results

### LAMP primer design and assay optimization

The four primer sets suggested by the Primer Design Tool from the conserved ITS2 region that contains unique *An. stephensi* sequence were tested against extracted DNA from three colony *An. stephensi* and eight other *Anopheles* species – *An. gambiae s*.*s, An. coluzzii, An. arabiensis, An. gambiae/coluzzii hybrid, An. funestus, An. quadriannulatus, An. dirus, and An. merus*. The two primer sets (P2L-45 and P26L2) that showed correct color change in *An. stephensi* and minimum cross-reactivity among other species were tested with varying concentrations and three incubation times (15 min, 30 min, and 45 min) to isolate the best candidate. No color change was detected at 15 minutes and specificity was affected at 45 minutes confirming 30 minutes as the ideal assay incubation time. A total of 384 reactions were analyzed in duplicate (total of 768 reactions) using eight different primer concentration combinations and primer set P26L2 was chosen for its consistent sensitivity and specificity (Table 2). The chosen primers and respective concentration combinations were tested using a temperature gradient (57°C, 61°C, 65°C, 69°C, 73°C, and 83°C) through 176 separate reactions. It was confirmed that primer concentration combination B3 at 65°C yielded the most consistent and specific results. Amplification (color change) still occurred at 69°C and 73°C but specificity results were inconsistent at these temperatures. No amplification was seen at higher or lower temperatures.

**Table 2.** Two primer set candidates were tested with differing primer concentrations to determine impact on assay sensitivity and specificity. Primer P26L2 at concentration B3 was found to produce optimal and consistent results.

| Conc.<br>combination | Primer concentration |  |  |  | Loop<br>Prime<br>r | Primer Set |  |  |  |
| --- | --- | --- | --- | --- | --- | --- | --- | --- | --- |
|  | F3 | B3 | FIP | B IP |  | P26L2 |  | P2L-45 |  |
|  |  |  |  |  |  | Sensitivit<br>y | Specificit<br>y | Sensitivit<br>y | Specificit<br>y |
| A1 | 2μM | 2μM | 16μM | 16μM | 4μM | X | ✓ | X | X |
| A2 | 4μM | 4μM | 16μM | 16μM | 4μM | ✓ | X | X | X |
| A3 | 2μM | 2μM | 32μM | 32μM | 4μM | ✓ | X | X | X |
| A4 | 2μM | 2μM | 16μM | 16μM | 8μM | X | ✓ | X | X |
| B1 | 5μM | 5μM | 40μM | 40μM | 10μM | ✓ | X | ✓ | X |
| B2 | 2.5μM | 2.5μM | 40μM | 40μM | 10μM | ✓ | X | X | ✓ |
| B3 | 5μM | 5μM | 20μM | 20μM | 10μM | ✓ | ✓ | X | X |
| B4 | 5μM | 5μM | 40μM | 40μM | 5μM | ✓ | X | X | X |

Repeat time-interval testing of 384 samples with the chosen primers showed no amplification (i.e. no color change) prior to 25 minutes, optimum amplification at 30 minutes, and a decrease of specificity after 35 minutes. Consequently, a 30-minute incubation was adopted for the assay. Once stopped (by being removed from the heat source), the product and color change remained stable and unaltered at room temperature for at least 12 weeks.

### CLASS assay analytical sensitivity

To test the sensitivity of the CLASS assay, a DNA serial dilution (1:10) of initial DNA extract concentration of 311.6 ng resulted in 100% positive color change to yellow. Yellow color change was repeatedly observed at 0.0003 ng concentration and greater; however, while concentrations lower than 0.0003 ng yielded positive color change, it occurred less consistently (33.3% of the time). Since these lower concentrations did not yield 100% color change, the assay sensitivity threshold established in this study is as low as 0.0003 ng, which is 1000 times lower than what is found in typical DNA extract from a single leg using our laboratory extraction method indicating high assay sensitivity (Table 3).

**Table 3.** The sensitivity of the CLASS assay was assessed using a serial (1: 10) dilution from DNA extract containing 311.6 ng of *An. stephensi* (UCI) whole mosquito DNA and resulted in 100% positive color change down to 0.0003 ng. Each dilution test was performed in triplicate. Color change at 1x10^7^ was inconsistent and samples with DNA concentrations lower than 0.0003 ng are less likely to result in a positive color change. and did not display as clear pink or yellow.

| Template dilution | Template concentration<br>(ng) | UCI |
| --- | --- | --- |
| NTC | - | Neg (3/3) |
| 1 | 311.6 | Pos (3/3) |
| 10 | 31.16 | Pos (3/3) |
| 1X10<sup>2</sup> | 3.12 | Pos (3/3) |
| 1X10<sup>3</sup> | 0.312 | Pos (3/3) |
| 1X10<sup>4</sup> | 0.031 | Pos (3/3) |
| 1X10<sup>5</sup> | 0.003 | Pos (3/3) |
| 1X10<sup>6</sup> | 0.0003 | Pos (3/3) |
| 1X10<sup>7</sup> | 0.00003 | Pos (1/3); Neg (2/3) |

### CLASS assay specificity and cross-reactivity

The optimized P26L2 primers (Table 1) were tested against extracted DNA from 11 laboratory-reared anopheline strains, including three *An. stephensi*. The assay was run 11 separate times with different extracted DNAs from single whole colony mosquitoes for a total of 132 reactions (Fig 2). Specificity was further assessed by testing DNA from 96 individual mosquitoes from each *An. stephensi* laboratory strain. One hundred percent of the samples yielded a positive result. Cross-reactivity was tested by sampling DNA from 48 *An. gambiae*, 48 *An. coluzzii*, 48 *An. arabiensis*, 48 *An. funestus*, and 48 *Ae. aegypti* whole mosquitoes and analyzed. None (0%) of the non- *An. stephensi* strains showed color change. All specificity assays were run in triplicate (Fig 2).

**Figure 2.**
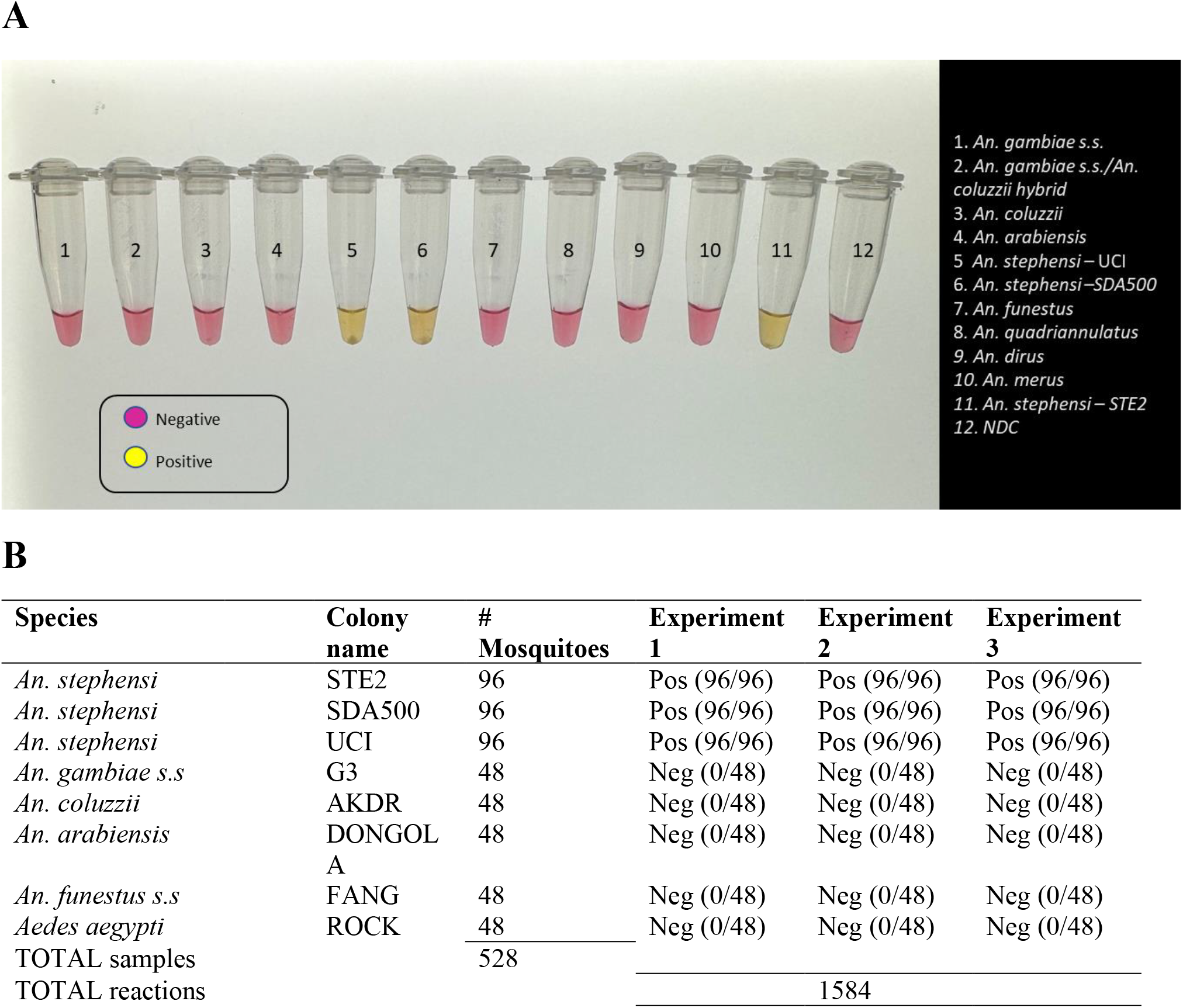
The *An. stephensi* CLASS assay shows 100% specificity when genomic DNA was tested in triplicate against 11 insectary and sequence confirmed mosquito strains, including three *An. stephensi* strains of different origins. **A**. Visualization of the CLASS assay shows positive samples resulting in a color change to yellow while negative and no template controls remain pink. NDC= No DNA template control. Samples were visualized on a white background and photographed on a standard light box. **B**. To determine specificity, the CLASS assay was run in triplicate on extracted DNA from 96 individual mosquitoes from each of the three *An. stephensi* insectary strains and DNA extracted from 48 individual mosquitoes from each species of *An*. gambiae s.s, *An. coluzzii, An. arabiensis, An. funestus*, and *Ae. aegypti*.

### CLASS assay testing of mosquito tissue, DNA extract, and DNA pooling

The assay was tested using a single insectary-reared mosquito leg inserted directly into the master mix and compared to DNA extracted from a single leg. Using DNA extract from a single leg resulted in amplification and color change was observed in *An. stephensi*, with no cross-reactivity with other tested species. Inserting a single mosquito leg straight into the master mix also successfully amplified following optimization, but at 35 min incubation time (Fig. 3), slightly longer than when using DNA extract. When testing a whole larva or whole adult mosquito into the master mix, the assay had low specificity, and yielded cross - reactivity; however, the use of DNA extract from a single larva or adult mosquito from 11 *Anopheles* colony strains, including three *An. stephensi*, and one *Ae. aegypti* resulted in positive color change (Fig 3). Limited testing on larval pan water yielded inconclusive results. Although the CLASS assay was able to identify *An. stephensi* from larval pan water and did not react with other Anopheline larval water source, it showed cross-reactivity with *Ae. aegypti*.

**Figure 3.**
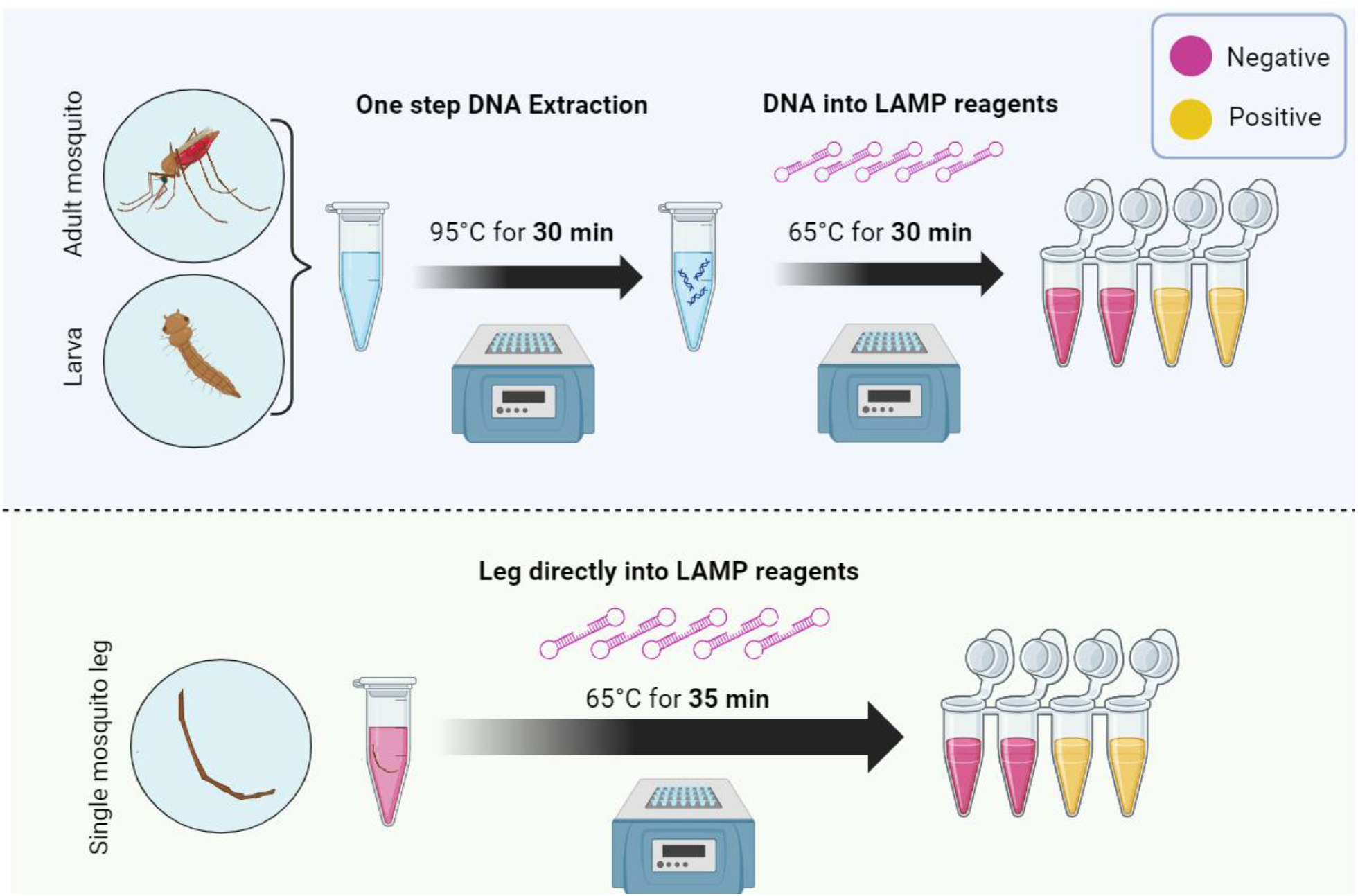
The CLASS assay shows high sensitivity and specificity when DNA extract from adult or larval mosquitoes is used and can also be used on a single mosquito leg directly with a 5-minute extension on incubation time, without the need for DNA extraction. DNA from any mosquito source, or a single leg directly placed in the colorimetric master mix are incubated at 65°C for 30 -35 minutes to obtain a yellow color change in a positive sample. Schematic produced using Biorender.com.

### CLASS assay specificity in field samples and comparison with conventional PCR method

To determine initial sensitivity on field-collected specimens, the assay was tested against three *An. stephensi* samples collected from Marsabit, Kenya (Ochomo et al., 2023); DNA from other cryptic species were also tested for cross-reactivity. The same samples were assayed using the conventional Singh PCR method for comparison. The assay showed a positive result in *An. stephensi* samples. There was no cross-reactivity across any other species tested. Additionally, DNA from 9 different colony-reared species *(An. gambiae ss, An. coluzzii, An. gambiae/coluzzii hybrid, An. arabiensis, An. funestus, An. quadriannulatus, An. merus, An. dirus*, and *Ae. aegypti)* and one *An. stephensi* (SDA 500) were combined to mimic a mosquito pool. The assay amplified as expected and was able to detect *An. stephensi* (Fig 4).

**Figure 4.**
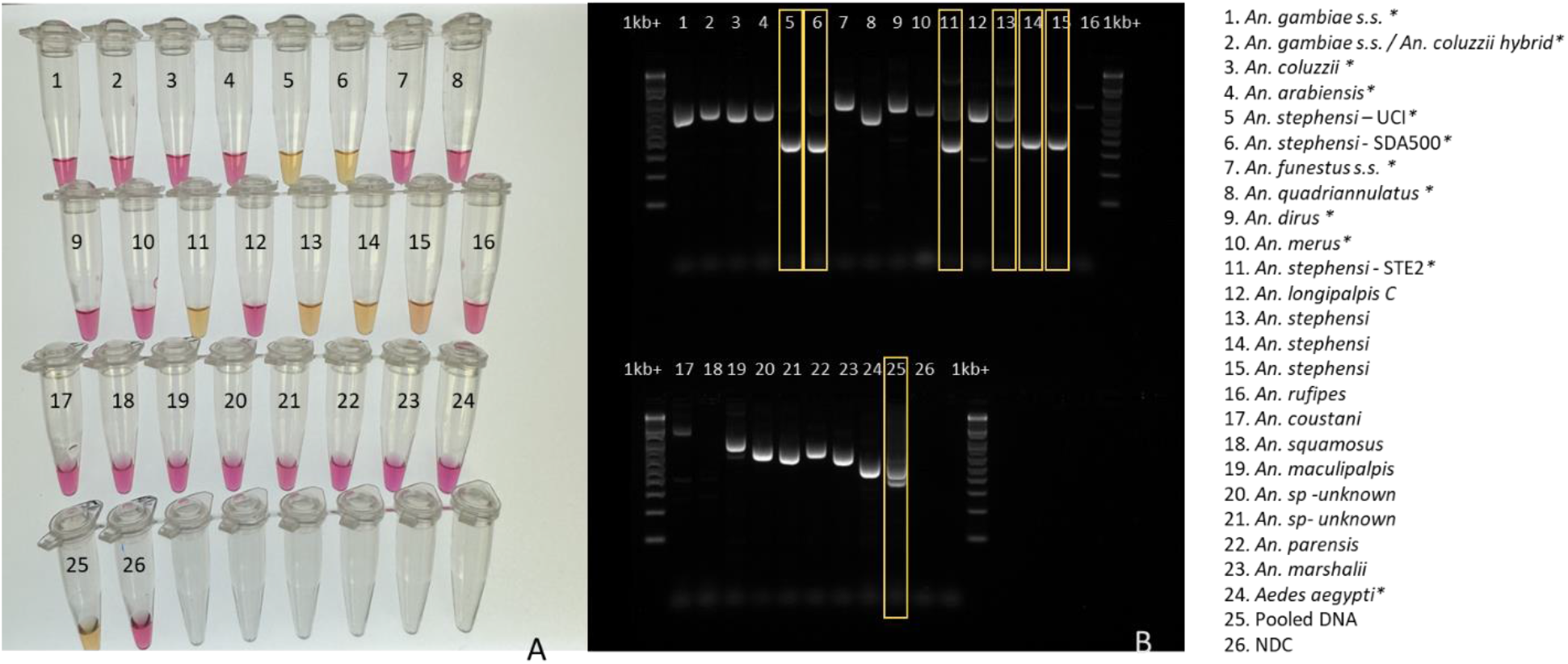
To validate the CLASS assay, comparison with the existing published species identification assay was conducted. Results from CLASS assay **(A)** *An. stephensi* 2023 Singh et al. PCR assay and **(B)** with assorted mosquito species is shown. Yellow boxes highlight *An. stephensi* products from *An. stephensi* PCR and the variation in results observed. Asterisk* samples 1-11, and 24 came from insectary –reared mosquitoes. Samples 12-23 came from sequence-confirmed field collected specimens. Sample 25 contained a pool of assorted mosquito DNA species, with *An. stephensi* represented 1:10. For both assays, 1µl extracted DNA was used.**Table 1**. Mosquito species used in this study and their corresponding strains and catalog numbers from BEI resources.

### CLASS assay testing of field samples from Marsabit, Kenya

CLASS assay testing of 55 COI sequence-confirmed samples collected in Marsabit, Kenya in 2023 successfully identified the 9 *An. stephensi* samples. Furthermore, no cross-reactivity was observed with other confirmed species or unknown samples (Table 4). The assay was duplicated to confirm results. There were 12 non-amplifications in the sample set during barcoding which also tested negative with the CLASS assay.

**Table 4.**
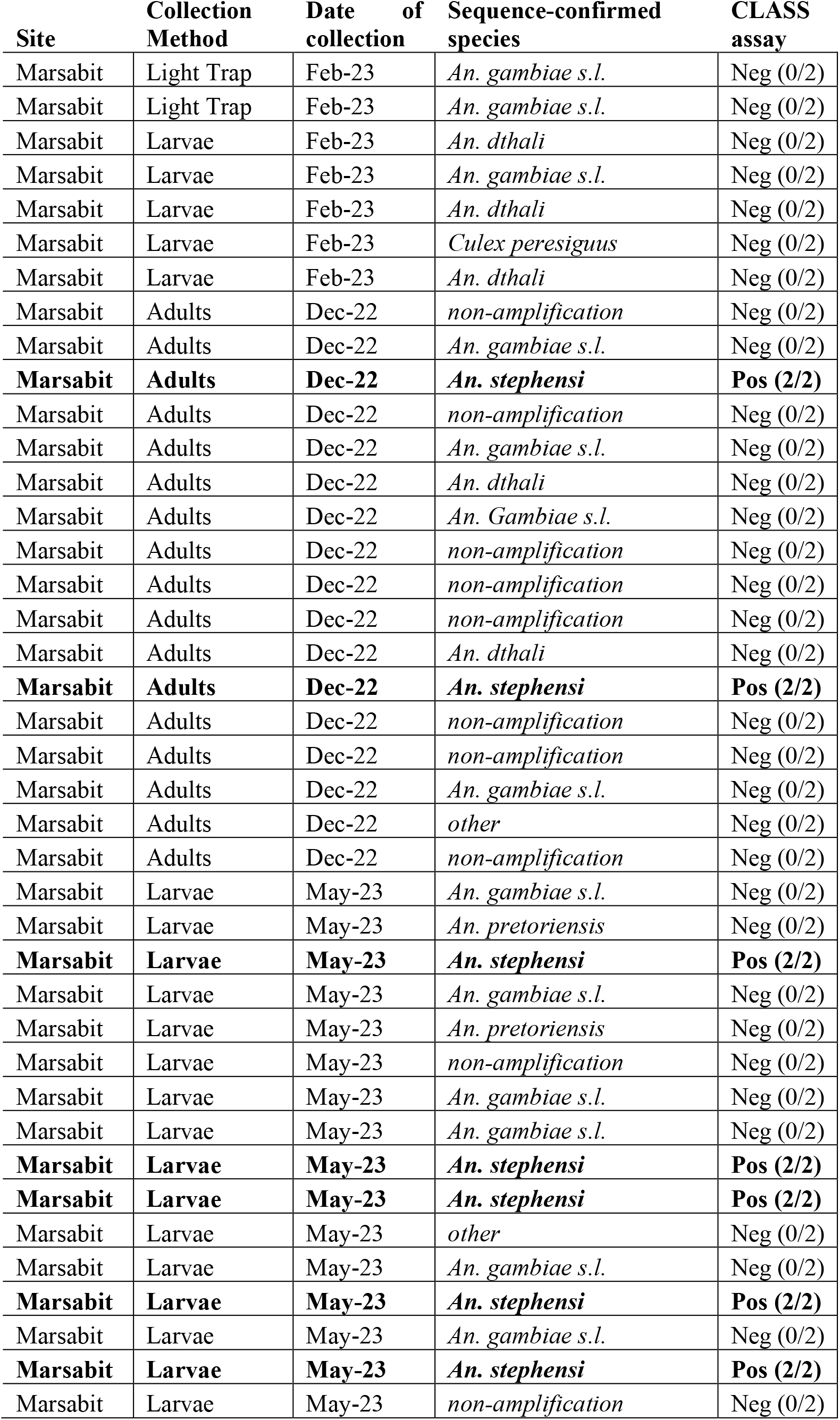
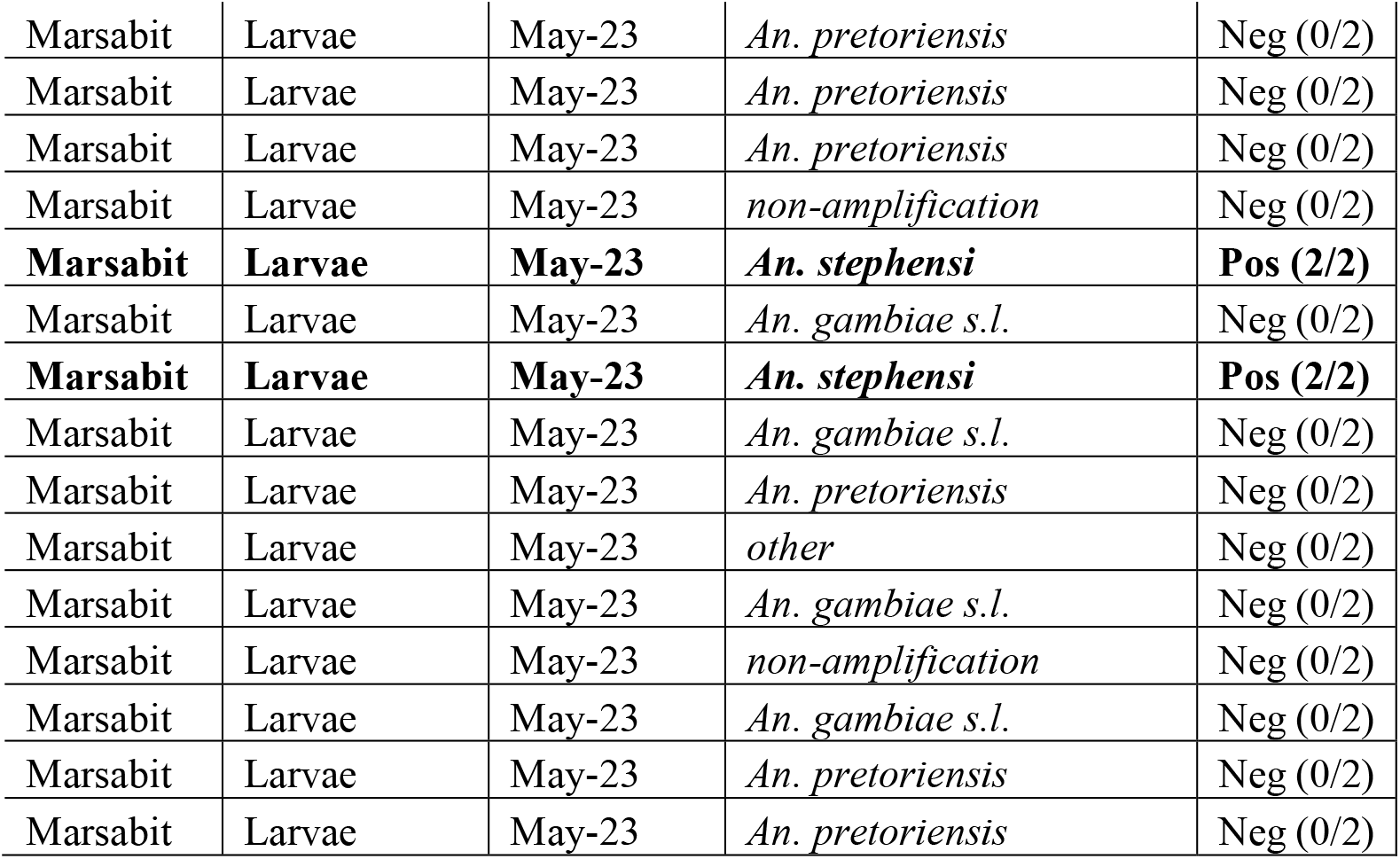
Wild *Anopheles* spp. collected in Marsabit, Kenya in 2022 and 2023 and which failed to amplify at KEMRI during routine species ID assays were sequenced using COI to confirm species identification. DNA extract from these samples was also run in duplicate through the CLASS assay. There is 100% concurrence between the CLASS assay results and Sanger sequencing in determining which specimens were *An. stephensi* and which were not.

## Discussion

Molecular species identification of malaria vectors is pivotal for effective control and elimination strategies which target species bionomics, particularly because malaria mosquitoes can often only be morphologically identified to the species complex or species group levels. Of increasingly complexity is the introduction of invasive species, like *An. stephensi*, which were not included in traditional identification keys (Gillies and Coetzee, 1987), but have now been added (Coetzee et al. 2020) yet can still be easily misidentified if morphological identification training is not conducted. Additionally, to confirm the presence of *An. stephensi* and document the spread of *An. stephensi*, surveillance reports of sites that are positive or negative are being added to the WHO *An. stephensi Threats Map* (WHO, 2023); however, due to lack of reliable molecular tools, at this point in time confirmation of the species in new countries requires molecular confirmation through Sanger sequencing (WHO, 2022) which results in delays in reporting.

A rapid one-step colorimetric LAMP assay was developed for species identification of *An. stephensi* to accelerate tracking of this invasive species across the African continent and other areas that have or are currently being invaded by this vector or where it is endemic. The CLASS identification assay provides a precise and reliable means of *An. stephensi* identification. The data presented here indicate high sensitivity and specificity of the assay whether mixed in a pool of 10 other species or validated against 8 species (including three unique insectary reared strains and wild caught invasive individuals of *An. stephensi*) and no false positives or false negatives have been reported yet. When a dilution series was conducted to determine analytical sensitivity, even at 0.0003 ng of DNA, the CLASS assay still detects the presence of *An. stephensi* DNA. Thus far, the specificity remains 100% when other species are processed through the assay. The ability to differentiate between various *Anopheles* and *Aedes* species, especially those with differing vectorial capacities or behaviors, is indispensable for tailoring interventions to specific vector populations (Sherrard-Smith et al., 2019).

The CLASS assay developed here can be run using a single leg from a suspect mosquito or DNA extract from an adult or larval mosquito (Fig. 3). DNA extracted from a wing or leg is preferable so voucher specimens can remain largely intact for further curation, sequencing, and storage as reference specimens. The use of an entire mosquito or larva for CLASS processing is highly discouraged as it yields non-specific results and prevents further follow up and species confirmation. For WHO submission and confirmation, sequencing positive specimens is still encouraged; however, the CLASS assay provides a rapid, high throughput, field friendly screening tool for an initial detection of *An. stephensi*. Although initial testing of larval pan water yielded inconclusive results, possibly due to Aedes excessive larval shedding in the water, all these findings suggest the potential for additional applications of the method. Future work could examine environmental DNA samples or large pools of specimens to confirm application and field deployment of the assay and would yield further information about potential cross-reactivity with other cryptic *Anopheles* species in natural settings.

In 2023 a conventional PCR assay was published (Singh et al. 2023) to support molecular detection of this species; however, conventional PCR may be a challenge in resource limited settings where molecular laboratory facilities and trained personnel are limited or absent. The Singh et al. (2023) assay has multiple primers and thus, there may be difficulty in interpreting positive results if one or both bands are absent. When running a number of insectary-reared and wild caught *Anopheles* species through the Singh et al. (2023) assay, the gel images showed a number of bands which could potentially be misinterpreted as false positive or false negatives. For example, *An. longipalpis* C, *Ae. aegypti*, and *An. coustani* produce bands and/or band sizes which could be misinterpreted as *An. stephensi*, and Sanger sequence confirmed *An. stephensi* from Kenya in this study inconsistently produce double bands in this PCR assay. Additional laboratories have also reported non-amplification of samples when using the Singh PCR assay, which when followed up by sequencing were later confirmed to be *An. stephensi* (Ochomo et al, 2023; Ochomo, personal communication, November 2023). As shown here, even insectary reared *An. stephensi* have the potential to only produce one band and hence inconclusive results using the existing PCR assay. Taken together, these findings indicate the need for a robust *An. stephensi* species identification assay that is simple to interpret and can handle rapid and high throughput processing of specimens. Follow up PCR and Sanger sequencing validation is still of critical importance, but for screening large numbers of samples a simpler approach is needed.

Results of the CLASS assay when testing field samples from Kenya show promising results. Although the testing of all field-collected samples was done using extracted DNA, its ability to use single legs without extraction, and its simpler equipment needs, gives the CLASS assay the potential for screening large numbers of samples in the field. With additional field deployment, enough data may be generated to potentially consider the CLASS assay as a species confirmation tool if its confirmed sensitivity and specificity continue to lie within at least an 85% confidence interval.

LAMP assay technology has substantially improved due to the need for rapid cost-effective diagnostics during the COVID-19 pandemic. With phenol-based colorimetric LAMP assays now widely adaptable, there are opportunities that exist beyond *An. stephensi* species identification. For example, *An. arabiensis*, a common malaria vector in Africa cannot be morphologically identified to the species level and therefore at the *An. gambiae* s.l. identification level requires an additional PCR assay (Scott et al., 1993) for species confirmation. With the colorimetric LAMP tools available, the existing rapid screening tool for *An. arabiensis* identification following *An. gambiae* s.l. morphological identification could be adapted and used in the field (Bonizzoni et al, 2009). In some locations, for example Ethiopia, *An. arabiensis* is the primary malaria vector and a screening tool like this could be used to identify *An. arabiensis* and distinguish non- *An. arabiensis* for further molecular confirmation. The original detection of *An. stephensi* in Ethiopia was found when specimens originally identified as *An. gambiae* s.l. were further examined molecularly through PCR, did not amplify as *An. arabiensis*, and sequencing revealed invasive *An. stephensi* (Carter et al. 2018).

Molecular species identification provides crucial data for epidemiological surveillance. By tracking the spread of *An. stephensi* and other invasive species, researchers and public health officials can monitor trends and adapt control strategies accordingly. Real-time data on vector distribution and density guide and the implementation of vector control methods, such as insecticide-treated bed nets and indoor residual spraying, ensure that resources are utilized effectively to curb malaria transmission (Chitnis et al., 2008). Early detection, facilitated by rapid assays like the CLASS assay, is critical for initiating timely responses to detections of invasive vector populations (Sinka et al., 2020).

The significance of accurate molecular identification of vector species extends beyond invasive *An. stephensi*. Long-term research and malaria program initiatives, guided by species identification data, enable program managers and scientists to study vector biology, behavior, and genetics. These insights are invaluable for developing effective control tools and strategies. Additionally, policy formulation relies heavily on accurate surveillance data. Molecular surveillance of vectors like *An. stephensi* informs policy decisions at regional, national, and international levels, ensuring a coordinated and effective response to malaria (Yakob & Yan, 2009) and accurate molecular identification not only aids in understanding the geographical distribution but also assists in predicting potential disease outbreaks, allowing public health authorities to proactively allocate resources and plan interventions (Dye, 1992).

## Conclusion

In conclusion, the molecular species identification of malaria vectors, particularly in the context of invasive species like *An. stephensi*, is indispensable to ensure gains made in global malaria control and elimination over the last few decades are not lost. The development of rapid, cost-effective assays such as the CLASS assay marks a significant advancement in the ability to detect this invasive malaria vector early in new locations for rapid response and its potential use as a screening tool to closely monitor the spread and establishment of the species into new locations. This tool is field adaptable and does not require a full molecular laboratory set up or highly trained molecular biologists for interpretation and therefore can be used in resource limited settings so that laboratory capacity is not a bottleneck preventing countries from detecting and reporting the species’ presence. The CLASS assay can not only facilitate early detection and accurate identification but can also be used to play a crucial role in understanding the evolving dynamics of malaria transmission by this invasive vector. By combining accurate molecular identification of *An. stephensi* with adaptive interventions, policymakers, researchers, and public health officials can work collaboratively to mitigate the impact of *An. stephensi*, to continue to work towards a malaria-free future globally.

## Supporting information

CLASS protocol supplement

## Disclaimer

The findings and conclusions expressed herein are those of the authors and do not necessarily represent the official position of the U.S. Centers for Disease Control and Prevention (CDC) or U.S. President’s Malaria Initiative (PMI).

## Acknowledgements

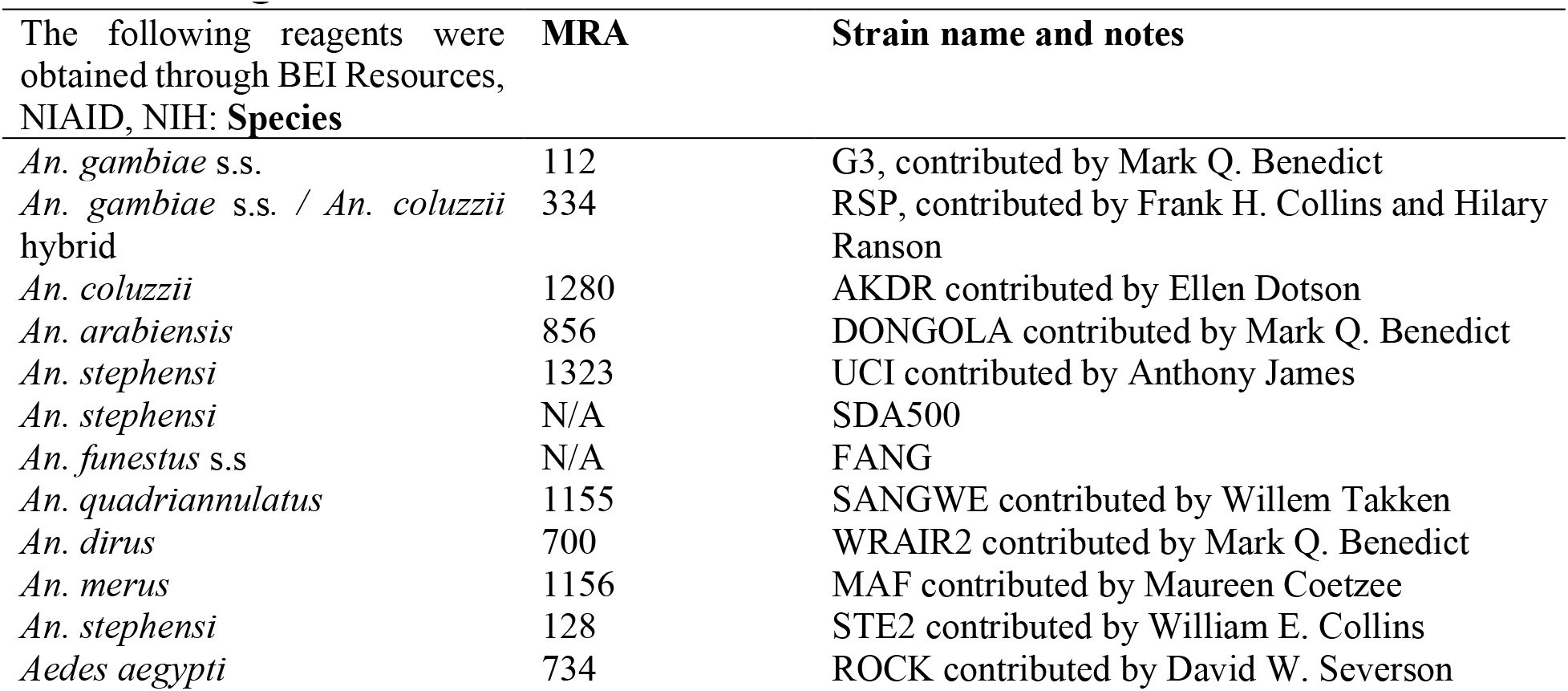

We thank the Malaria Research and Reference Reagent Resource Center (MR4) staff at the US Center’s for Disease Control and Prevention (CDC) Entomology Branch, particularly Laura Leite and Catherine Steele for their help and support rearing and providing colony mosquito material for this study. We would like to thank Biorender.com for the support in generating figures for the study. Gloria Raise and JeNiyah Scaife were funded through Public Health Entomology for All (PHEFA), a partnership with the Entomological Society of America (ESA) and the US CDC. Financial support for CR and SZ was provided by the US President’s Malaria Initiative (PMI). We thank Anne Powers, Holley Hooks, the Division of Vector Borne Diseases (DVBD) and the National Center for Emerging and Zoonotic Infectious Diseases (NCEZID) for the PHEFA program. We also thank John Gimnig, Ellen Dotson, Audrey Lenhart, Lisa Reimer, and the Entomology Branch for thoughtful and crucial feedback on the concept and assay development.

