## Supplementary material for "A rapid, cost-effective, colorimetric LAMP assay (CLASS) for detecting invasive malaria vector, *Anopheles stephensi*": CLASS protocol supplement

### **Supplementary File**

**Colorimetric LAMP *Anopheles stephensi* identification assay (CLASS) protocol**

This assay utilizes Loop Mediated Isothermal Amplification (LAMP) and a phenol-based color change to quickly screen and identify *An. stephensi*.

1.0 **Procedure/Guidelines**
    1.1 Prepare MM for reactions as follows:

| **Volumes (µl) per Reaction Number** | | | **Component** | **Working Concentration** | **Primer Sequence**  **(Rafferty et al, 2024)** |
| --- | --- | --- | --- | --- | --- |
| **1** | **50** | **100** |  |  |  |
| 9 | 450 | 900 | Deionized pure water | **___** | **_____** |
| 12.5 | 625 | 1250 | WarmStart^®^Colorimetric LAMP 2X Master Mix, | 5X | **_____** |
| 0.5 | 25 | 50 | F3 | 10µM | ATTGCACGGGGACTTCCA |
| 0.5 | 25 | 50 | B3 | 10µM | GCCTACAGACTCCACTGTCA |
| 0.5 | 25 | 50 | FIP | 40µM | CGACTGCAACTGTATGCGAGGACGGGTCGAGTAACACTTGC |
| 0.5 | 25 | 50 | BIP | 40µM | CCGTGTGGGTGAGTGAGGTTAGAATGATGCGACGGGAGAAG |
| 0.5 | 25 | 50 | LF | 20µM | AAGATACGAGCGCGTTGGG |
| **24** | **1200** | **2400** | **Total (to each 24 µl reaction add 1 µl template DNA) or see note 3.2** | | |

1.2 Incubate in a heat block or thermal cycler at 65 °C for 30 minutes.
1.3 Analyze samples visually against a white background for color change.

2.0 **Interpretation**
    2.1 A sample is considered positive when the color changes from bright pink to bright yellow.
    2.2 Samples with a light pink or orange color are considered negative but can be followed up for further confirmation.
3.0 **Notes**
    3.1 All assays should include a positive control (confirmed *An. stephensi* DNA), a negative control (confirmed DNA from other *Anopheles* or *Aedes* mosquito), and a no DNA control to ensure results are valid.
    3.2 A single leg can be used in lieu of DNA. If that is the case, add 25 µL of Mastermix to reaction, and incubate for 35 minutes.
    3.3 To conserve reagents, it is possible to halve the reaction by using the following guidelines:
        3.3.1. Ensure primers are at 10X concentration in the reaction.
        3.3.2. Do not use single legs or un-extracted mosquito tissue as it will yield false positives.
    3.4 Positive samples should be followed up with Singh (2023) assay and confirmed by sequencing.
